# Trapped in the web: network architectures spread coevolution and shape adaptation

**DOI:** 10.1101/2025.11.04.686579

**Authors:** Alexandre Fuster-Calvo, Cecilia S. Andreazzi, Christine Parent, François Massol, Paulo R. Guimarães, Dominique Gravel

## Abstract

Adaptation is critical for biodiversity to persist under global change. Within ecological communities, species often face trade-offs between adapting to shifting abiotic conditions and navigating the complex selective pressures imposed by interaction networks. We hypothesize that network architectures characterized by high interaction diversity and overlap constrain coevolutionary dynamics, with asymmetric outcomes for exploiters and victims. Specifically, we predict that exploiters, subject to spread and conflicting selection imposed by their victims, will evolve more slowly and show reduced capacity to track victims’ evolutionary responses, with these constraints strongest for generalist exploiters. In contrast, victims will show more variable dynamics depending on the coherence of selection (i.e., whether pressures from different exploiters push the victim’s trait in the same vs. different directions). To test this, we simulated trait evolution in coevolving communities of exploiters and victims across 91 empirical networks, and in artificial networks designed to isolate specific structural effects. Our results show that higher connectance, species richness, nestedness, and centrality homogenize biotic effects and increase fluctuations in trait matching, ultimately weakening coevolutionary coupling. Under these conditions, exploiters face conflicting selection that slows evolution, whereas victims either benefit from aligned selection that accelerates evolution or are constrained by multiple pressures. Together, our findings suggest that network architecture plays a fundamental role in shaping coevolution and adaptation, and raises broader questions about its influence on eco-evolutionary processes in more complex and environmentally variable systems.

## Introduction

Evolution contributes to biodiversity response to global changes (Norberg et al. 2012, Urban et al. 2016, 2024, Catullo et al. 2019, Carrol et al. 2023), often on ecological timescales (Post & Palkovacs 2009, Fronhofer et al. 2023, Jarne & Pinay 2023). Traits such as phenology (e.g., Vitasse et al. 2018), thermal tolerance (e.g., Geerts et al. 2015), responses to pollution (e.g., Whitehead et al. 2017), dispersal abilities (e.g., Ochocki & Miller 2017), and species interactions (Moran & Alexander 2014, Galetti et al. 2013) can evolve rapidly, generating eco-evolutionary feedbacks that alter community dynamics (Hendry 2017, Govaert et al. 2019).

While coevolutionary theory is well developed for pairwise interactions and small modules (Nuismer & Doebeli 2004, Thompson 2005), extending it to the complex interaction networks of natural communities remains a major challenge (Fussmann et al. 2007, Pillai et al. 2012, Barraclough 2015, Toju et al. 2017, Govaert et al. 2019, Calcagno et al. 2023, Cosmo et al. 2023). Foundational concepts like arm race escalation or Red Queen dynamics (Buckingham & Ashby 2022), developed for isolated species pairs, do not clearly explain emergent properties of species-rich systems. In these systems, direct and indirect effects generate coevolutionary cascades that reshape selection pressures across the network (Guimarães et al. 2011, McPeek 2017, Guimarães et al. 2017, terHorst et al. 2018, Hall et al. 2020). As interactions multiply, multiple partners become sources of selection, potentially generating spread and conflicting selection (Fox 1988, Johnson & Stinchcombe 2007), underscoring the need to account for network architecture to understand how coevolution unfolds in complex communities (de Meester et al. 2019, Hendry 2019) and anticipate responses to environmental change (Norberg et al. 2012, Loeuille 2019, Åkesson et al. 2021).

Modeling efforts have significantly advanced our understanding of coevolutionary dynamics in species-rich systems (e.g., Melián et al. 2011, Moya-Laraño et al. 2012, Levine et al. 2017, Toju et al. 2017, Hui et al. 2018, Patel et al. 2018). However, despite important advances, our understanding of how interaction network structure influences coevolutionary dynamics remains limited. While ecologists have explored in depth how network architecture facilitates or impedes the propagation of environmental perturbations and affects ecological dynamics (Boccaletti et al. 2006, Martins et al. 2024), less attention has been devoted to how it shapes evolutionary dynamics and the capacity of species to adapt to environmental change (Andreazzi et al. 2017).

In coevolving communities, a higher density of connections could propagate evolutionary effects more broadly but weaken their strength, spreading the consequences of selection. This outcome assumes that a species’ total interaction strength is constrained, i.e., having more partners means interacting less strongly with each of them, which should loosen trait coevolution between species (Cosmo et al. 2023). Conversely, more compartmentalized or modular networks could foster more direct and stronger interactions and selection within specific modules, leading to localized strengthening of coevolution (Guimarães et al. 2011, Andreazzi et al. 2017, Maia & Guimarães 2024). Multiple interaction partners introduce conflicting selection pressures, akin to particles in fluid systems: while particles move freely with few interactions, friction and opposing forces slow them down when connections increase. Similarly, species with a high degree—those interacting with many others and potentially occupying structurally important positions (Delmas et al. 2019)— may evolve more slowly, as their traits are shaped by multiple and opposing pressures.

In mutualistic networks, it has been shown that indirect effects conflict with adaptation to partners (Cosmo et al. 2023). In antagonistic networks, these effects are likely asymmetric, reflecting the contrasting effects of biotic selection experienced by exploiters and victims (Nuismer & Thompson 2006). In antagonistic interactions, victims such as prey or hosts may benefit from the slower evolutionary responses of their enemies, escaping more rapidly through faster adaptation. Multiple and conflicting selective pressures on exploiters can hinder their ability to track and counter evolving defenses, potentially weakening coevolutionary coupling between the two sides.

Of course, this is a simplified picture; real systems are more complex (Kopp & Matuszewski 2014). Trait responses depend not only on selection strength, but also on the source of adaptive variation—adaptation from standing genetic variation versus new mutations (Barrett & Schluter 2008); the amount and alignment of genetic variance across traits, including trait correlations that channel or constrain responses (Lande & Arnold 1983, Assis et al. 2020); and phenotypic plasticity, which can buffer or amplify short-term change (Merilä & Hendry 2014). Yet, we posit that in many antagonistic systems, more densely connected network structures lead to the spreading of selection and a slowdown in trait evolution, constraining adaptation in some species’ groups, while facilitating escape in others.

In this study, we focus on interaction network properties that influence the distribution and overlap of interactions across communities. We hypothesize that higher connectance, nestedness, and species richness, combined with lower modularity, should increase the potential for indirect effects and generate multiple and sometimes conflicting selection pressures (Guimarães et al. 2017). These effects should be especially pronounced for central species. Together, these network features are expected to shape the strength and coherence of selection across the community, influencing both the direction and pace of trait evolution. We investigate this proposition with simulations of trait evolutionary dynamics applied to artificial and empirical network structures. Our approach involves (1) quantifying how network features influence the spread of biotic interactions, and (2) investigating their effects on evolutionary rates, coevolution, and adaptation.

## Methods

We modeled species coevolution in antagonistic bipartite networks using discrete-time trait evolution frameworks developed by Andreazzi et al. (2017, 2020) and Guimarães et al. (2017). These models extend foundational approaches to phenotypic evolution (Lande 1976) by incorporating the reciprocal selective pressures generated through species interactions. Our objective is to detect relationships between network structure and species’ centrality and their ensuing impacts on evolutionary rates, coevolutionary dynamics, and adaptation to both biotic and abiotic environments. To do this, we fix the network structure throughout simulations (i.e., there is no interaction rewiring of the original topology). Furthermore, because isolating the effects of specific aspects of network structure is challenging, we complement our analyses with simulated networks. This approach enables us to manipulate specific metrics while holding others constant, providing deeper insights into their respective influences.

### Model

Biotic selection is formalized using a match-mismatch model (Nuismer & Thompson 2006, Nuismer et al. 2007, Hanifin et al. 2008, Yoder et al. 2010, Boots et al. 2014), a common formalization for trait-based models of Red Queen dynamics. This approach assumes that natural selection favors exploiters whose attack traits align with the defense traits of victims, increasing their likelihood of a successful attack. Conversely, selection favors victims whose defense traits diverge from those of exploiters, enhancing their ability to evade attacks.

Initially, we assign a real number *z*(0), randomly sampled from a normal distribution *N*(0,0.1), to represent the initial mean trait value of each species at the start of the simulation. These trait values then evolve over time in response to selection pressures. We assume that abiotic selection favors a fixed optimum trait value, 𝜃_*i*_, which we set equal to its initial trait value for simplicity. The partial selection differential caused by abiotic selection is defined as:

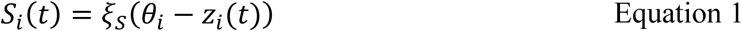

where 𝜉_𝑠_ is the intensity of abiotic selection on all species, and equals 0.5 throughout.

Biotic selection depends on the degree of trait matching between an interacting pair of exploiter *i* and victim 𝑗 species. Selection on exploiters favors matching their traits to those of their victims, while victims are selected against having traits matching those of their exploiters. For convenience, we define 𝐼_*i*_as the set of species interacting with exploiter *i*. The partial selection differential on exploiter traits, from victim 𝑗 𝜖 𝐼_*i*_, is:

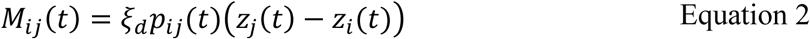

where 𝜉_𝑑_ is the selection intensity imposed by interacting victims, set to 0.5. This selection is weighted by the relative degree of matching between species pairs, 𝑝_*i*𝑗_(𝑡), which depends on phenotypic matching and the exploiter trait-based interaction sensitivity, 𝑏:

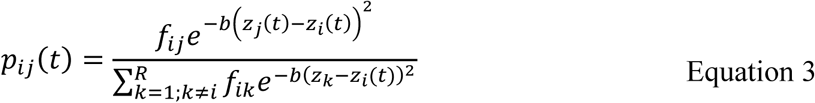

Where 𝑓_*i*𝑗_ is an element of the binary adjacency matrix 𝐹. This matrix specifies whether a potential interaction between species *i* and 𝑗 can occur (𝑓_*i*𝑗_ = 1) or not (𝑓_*i*𝑗_ = 0, i.e., a forbidden link; Olesen et al. 2011). 𝐹 is a square, species-by-species matrix with two off- diagonal blocks describing cross-guild interactions (victim-exploiter and exploiter-victim) and zeros in the diagonal blocks, since species do not interact within the same guild. 𝑅 is the total number of species in the network.

For victim species 𝑗, we define 𝐼_𝑗_ as the set of all exploiters interacting with 𝑗, and 𝐼_*i*_[*z*_𝑗_ − 𝜀, *z*_𝑗_ + 𝜀] as the subset whose trait mismatch Δ*z*_*i*𝑗_ does not exceed the threshold 𝜀 (set to 0.5 for all species). Only exploiters in this subset contribute to the selection gradient on 𝑗; those with Δ*z*_*i*𝑗_ > 𝜀 exert negligible biotic selection (Andreazzi et al. 2017). The partial selection differential on victim traits, from exploiter *i* 𝜖 𝐼_𝑗_[*z*_𝑗_ − 𝜀, *z*_𝑗_ + 𝜀], is:

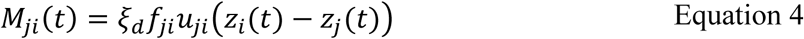

where:

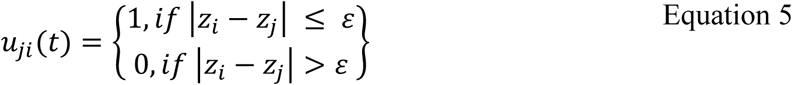

Selection favors moving the victim’s trait value away from the exploiter’s trait. If *z*_𝑗_(𝑡) > *z*_*i*_(𝑡), selection acts to increase *z*_𝑗_(𝑡) towards *z*_𝑗_(𝑡) + 𝜀; if *z*_𝑗_(𝑡) < *z*_*i*_(𝑡), selection acts to decrease *z*_𝑗_(𝑡) towards *z*_𝑗_(𝑡) − 𝜀. Beyond the threshold 𝜀, however, further divergence does not attract the victim toward any specific value, as the exploiter’s effect becomes negligible, and selection on *z*_𝑗_(𝑡) from this interaction is effectively neutral.

Combining equations 4.1 to 4.4 with the breeder’s equation results in a general formulation describing trait evolution for both exploiters and victims:

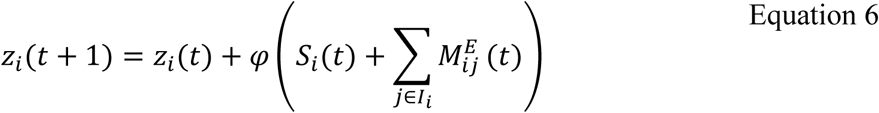

If *i* is an exploiter, and

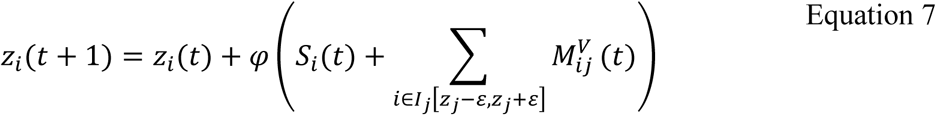

Where 𝜑 is a composite parameter that scales the selection gradient into evolutionary change and is proportional to the additive genetic variance and the slope of the fitness landscape (Guimaraes et al. 2017). 𝑆_*i*_(𝑡) is the partial selection differential due to environmental selection, and the sum over 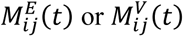 represents the total biotic selection imposed by interaction partners, depending on whether *i* is an exploiter or victim.

### Characterizing trait evolutionary responses

We calculate the evolutionary rate for each species as the average absolute change in its trait value across a specified time period 𝑇, starting at timestep 𝜏:

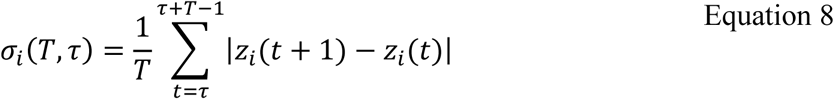

For community-wide evolutionary rates, we average the evolutionary rates across all species in the community:

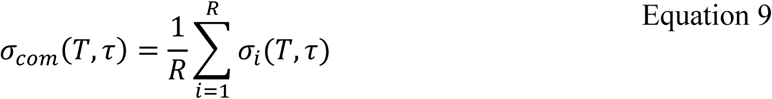

Where *R* is the number of species in the network. We quantify the fluctuation in trait matching for a given victim-exploiter pair over a specified time period 𝑇, starting at timestep 𝜏, as:

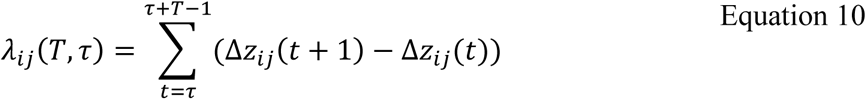

The mean temporal fluctuation in exploiter-victim trait differences over a period 𝑇 is calculated as:

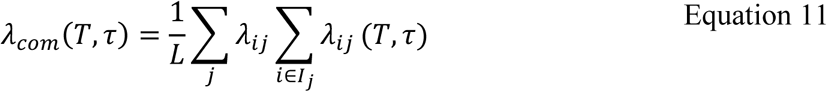

where 𝐿 is the number of interacting pairs (*i*, 𝑗) with 𝑓_*i*𝑗_ = 1. A high 𝜆 value indicates high temporal variation in interaction strength among species, which is a proxy of interaction fluctuating selection (de Andreazzi et al. 2017).

### Measuring adaptation to biotic interactions

We focus on two components of fitness load to investigate how network structure affects adaptation: biotic and network loads. These components reflect how well species traits align with those of their interaction partners and how structural properties of the network contribute to maladaptation. The total fitness load (or maladaptation) due to interactions quantifies the mismatch between a species’ trait and the traits of its interacting partners. For an exploiter *i*, it is defined as:

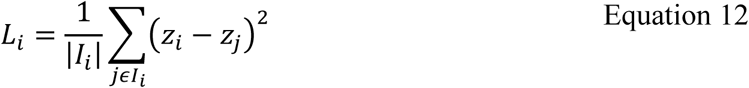

This formulation captures the deviation of the exploiter’s trait *z*_*i*_ from the traits of its interacting victims. To minimize this load, the exploiter’s trait value *z*_*i*_should converge toward an optimal value *z̄*_*i*_, its “biotic optimum”, given by:

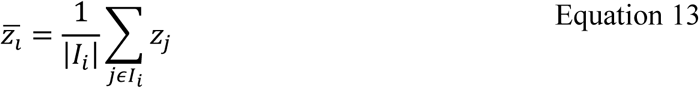

Using this biotic optimum, we can decompose the load into two distinct components. The biotic load (𝐿^𝑏*i*𝑜𝑡*i*𝑐^) quantifies the mismatch between the exploiter’s trait and the biotic optimum:

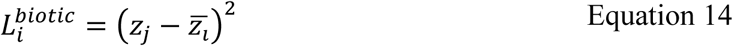

The second component, the network load (𝐿^𝑛𝑒𝑡𝑤𝑜𝑟𝑘^), is an incompressible load that is independent of the focal species trait value. It measures how much maladaptation is due to the variance of trait values among species in 𝐼_*i*_:

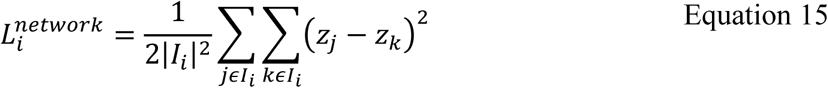

where 𝑗 and 𝑘 index all species in 𝐼_*i*_. The network load reflects the inherent constraints imposed by the structure of the network, as it cannot be mitigated by adjustments to the exploiter’s trait.

For the victim species 𝑗, the biotic load is defined differently due to their distinct selection regime, focusing on mismatches that allow them to escape exploiters:

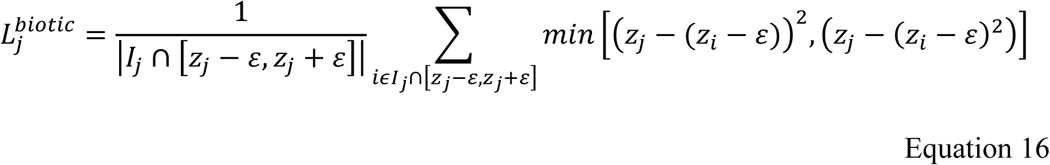

This term reflects the victim’s ability to evade exploiters. As defined above, beyond the threshold 𝜀, exploiters have negligible effects on the victim’s fitness, and the selection from these interactions becomes neutral. Consequently, victims’ adaptation focuses on escaping individual exploiters rather than responding to the variance among their traits, which is why we do not compute the network load for victims.

### Measuring biotic effects’ evenness

We investigate how network architecture influences the spread of biotic selective pressures among species. We analyze the total effects of biotic interactions within the network, incorporating both direct and indirect effects (Guimarães et al. 2017). We start by rescaling 𝐹 into a row-normalized form, 𝐹’, so that each row sums to one. This normalization ensures that the influence of one species on another is expressed as a relative contribution, making the effects comparable across different network structures:

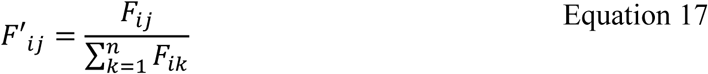

The normalized matrix 𝐹’ is then scaled by a scalar 𝑚, which specifies the total strength of biotic effects on a species in the same units as the selection coefficient in our evolutionary model. In our simulations, 𝑚 is set to 𝜉_𝑑_ = 0.5 for all species, but in principle, 𝑚 can be a diagonal matrix to allow species-specific strengths. This step results in the weighted matrix 𝑄:

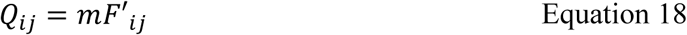

To compute the total effects (direct + indirect) among species, we calculate the matrix 𝐸 by solving the equation:

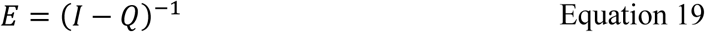

where 𝐼 is the identity matrix of the same dimensions as 𝑄. The elements of 𝐸 describe the total effect of species 𝑗 on species *i*, including both direct and indirect contributions through the network. To quantify the evenness of these total effects—“biotic effects’ evenness (H)”— we normalize each row of *E* to obtain *E’*:

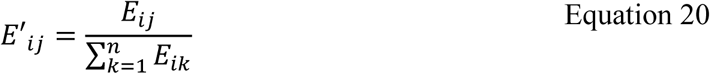

We then compute the Shannon index:

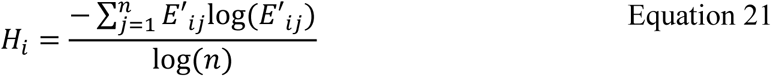

where *n* is the number of species. This normalization bonds 𝐻_*i*_ between 0 and 1, where 0 represents complete unevenness (all effects concentrated in one species) and 1 represents maximum evenness (effects equally distributed across species). A high value of 𝐻_*i*_ suggests that the effects of species on *i* are evenly distributed, meaning that each species contributes relatively equally to the overall impact on *i*. On the other hand, a low value of 𝐻_*i*_ indicates that the effects are unevenly distributed, with certain species exerting a disproportionately large influence compared to others. In addition to the normalized 𝐻, we also compute 𝐻 using raw values of the total effects matrix 𝐸 (i.e., without normalizing each row to sum to 1) to assess whether normalization significantly alters the results. To test this, we investigate the relationships between raw and normalized 𝐻 and network metrics, and calculate Pearson correlations between normalized and raw 𝐻 (Figure S3). This approach allows us to ensure that normalization does not introduce substantial deviations while maintaining consistency across networks of varying sizes.

### Empirical network structures

We use the 92 bipartite antagonistic networks used in de Andreazzi et al. 2017 to parametrize the model, exploring the effects of network topology on coevolutionary dynamics. These consist of 43 predation (37 bacteria-phage and 6 predator-prey), 31 host-parasite, and 18 herbivory networks. The networks range from small networks of 9 species of bacteria-phages to large networks of almost 308 species of leaf-mining herbivores and their host plants (Table S1, Figure S1).

We describe network topology by combining both the size of the network (i.e., number of species, *S*) and the density (i.e., connectance *C*, defined as the proportion of realized links between species relative to the total number of possible links, *C*), using the complexity index √𝐶𝑆 (May 2001, Jacquet et al. 2016). This index encapsulates how “densely packed” the network is with interactions, considering both the number of species and their propensity to interact. At the species level, we measure species’ degree—the number of distinct partners a species interacts with (Freeman 1977).

### Simulated network structures

We simulated bipartite networks in two main steps. First, we defined the expected number of partners (degree sequence) for each species in both guilds, setting these values to match the structural property we wanted to vary (species richness, connectance, degree distribution, or nestedness). Second, we generated a binary interaction matrix from these expected degrees using a Bipartite Expected Degree Distribution (BEDD) model (Ouadah et al. 2022), which produces networks that match the target degrees and connectance on average. In Step 1, we varied one structural property at a time while keeping the others fixed at their empirical averages. Following the approach described by Massol et al. (2021), the number of partners for victims and exploiters was represented as two vectors, 𝐷_1_and 𝐷_2_, where 𝑛_1_ and 𝑛_2_ are the numbers of victim and exploiter species, respectively. In all cases, the resulting 𝐷_1_ and 𝐷_2_ defined the target degree sequence for step 2.

When varying richness, total richness S = 𝑛_1_ + 𝑛_2_ ranged from 12 to 120 in steps of two, with the ratio 𝑛_1_⁄𝑆 and connectance *C* fixed to the empirical mean. Degrees were then set to 𝐷_1,*i*_ = 𝐶𝑛_2_ and 𝐷_2,𝑗_ = 𝐶𝑛_1_ for all exploiters *i* and victims 𝑗, yielding a Poisson-like degree distribution. When varying connectance, *C* ranged from 0.05 to 0.90 in steps of 0.01, with 𝑛_1_ and 𝑛_2_ fixed to their empirical means, and the same Poisson-like degree family was used. For the nestedness gradient, by contrast, we used a constructive nested degree generator in which each victim has exactly 𝑛_𝑛𝑒𝑠𝑡_ expected partners (chosen at random columns), with 𝑛_𝑛𝑒𝑠𝑡_ ranging from 1 to 29 in increments of 0.5. This procedure inherently changes the degree distribution and allows connectance to vary naturally, producing increasing NODF (Nestedness metric based on Overlap and Decreasing Fill; Almeida-Neto et al. 2008) values. Because this structural change also influences modularity (Payrató-Borràs et al. 2019), we quantify modularity using four different algorithms – given that each of them can yield slightly varied results depending on the network’s characteristics (Oza et al. 2023): the Louvain method (Blondel et al. 2008), the Walktrap algorithm (Pons & Latapy 2005), the Fast-Greedy algorithm (Clauset et al. 2004), and the Leading Eigenvector method (Newman 2006). For each network, the modularity value is calculated as the mean of these four measures.

We explored two contrasting degree-distribution families that represent opposite extremes of network architecture. In scale-free networks, degrees are highly heterogeneous, with many specialists and a few generalists forming heavy-tailed degree distributions (Barabási & Bonabeau 2003). In contrast, core-periphery networks have strongly ordered degrees: a densely connected “core” of generalists and a sparsely connected “periphery” of specialists, producing high nestedness (Martín González et al. 2020). Here, the degree sequence is not simply a function of 𝑛_1_, 𝑛_2_, and *C* as in the other treatments, but requires a specific statistical form. In scale-free networks, the degree sequence follows a heavy-tailed distribution, with a few highly connected hubs and many nodes of low degree, but without an imposed ordering of interactions. We generated these sequences using the Barabási-Albert preferential-attachment algorithm (Barabási & Albert, 1999; as implemented in *igraph*), in which nodes are added sequentially and linked to existing nodes with probability proportional to their current degree, yielding power-law-tailed 𝐷_1_and 𝐷_2_. The number of nodes in each guild is set to match the empirical averages, and the parameter 𝑚, which controls the number of edges added per new node, was set to *m* = 1. To maximize the contrast with the scale-free case, we simulated the most nested core-periphery networks possible for the given 𝑛_1_ and 𝑛_2_. Degrees were perfectly ordered in each guild: the most generalist victim connected to all exploiters, the next to all but one, and so on, with exploiter degrees assigned in the same ordered fashion. This produces the steepest possible ordered degree distribution, yielding maximal nestedness. In our implementation, this corresponds to setting the hierarchical assignment parameter *N*_𝑠_to a high value (*N*_𝑠_= 12), where *N*_𝑠_ defines the number of exploiters connected to the most specialized victim, with other victims assigned degrees decreasing linearly from 𝑛_2_ (all exploiters) down to *N*_𝑠_.

In step 2, we used the target degree sequences from Step 1 to generate a binary biadjacency matrix A using the BEDD model. For each victim-exploiter pair, we computed the connection probability:

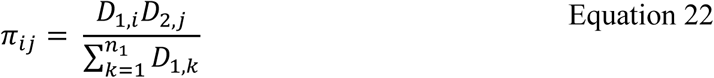

where 𝐷_1,*i*_is the target degree of exploiter *i*, 𝐷_2,𝑗_ is the target degree of victim 𝑗, and the denominator sums over all 𝑛_1_exploiters. The biadjacency matrix *A* has entries 𝐴_*i*𝑗_ indicating the presence (1) or absence (0) of an interaction between *i* and 𝑗, and each 𝐴_*i*𝑗_is drawn independently from a Bernoulli distribution with parameter 𝜋_*i*𝑗_. This procedure ensures that the expected degrees of the realized network match the target sequences, and that global properties such as connectance or nestedness match the values implied by those degrees on average. When the sampling produces isolated species, we add a single link to each isolate, preferentially attaching it to the most connected partner, to avoid disconnected nodes. Across all treatments, this design yielded 398 simulated networks: 200 varying in degree-distribution family (100 scale-free, 100 highly nested), 86 varying in, 57 varying in nestedness, and 55 varying in species richness.

### Simulations

Simulations are conducted in stable, unperturbed environments, where species’ environmental optima 𝜃_*i*_ are held constant. The system is allowed to evolve until equilibrium, defined as 20 consecutive timesteps without changes in trait values for all species within a tolerance of 10^−10^. At equilibrium, the trait values of all species are recorded. We evaluate trait evolutionary rates, fluctuations in trait matching, and levels of adaptation for both victims and exploiters. The analysis is based on 20 replicates of each of the 92 empirical networks and 15 replicates of each of the 398 simulated networks.

## Results

Exploiters and victims respond oppositely to network structure in terms of evolutionary rates. Simulations with synthetic networks show that species richness, connectance, and nestedness reduce exploiters’ community-averaged evolutionary rates, while modularity increases them (Figure 1A,D,G,J). In contrast, victims display faster evolutionary rates with increasing richness, connectance, and nestedness and slower rates with increasing modularity (Figure 1B,E,H,K). These observations are mostly consistent with those from empirical networks: increases in complexity index are associated with slower evolutionary rates in exploiters and faster rates in victims (Figure 2A,C,D,F). In contrast, nestedness and modularity show no consistent effects, except for a tendency toward faster victim evolution in more modular networks (Figure 2B,E).

**Figure 1.**
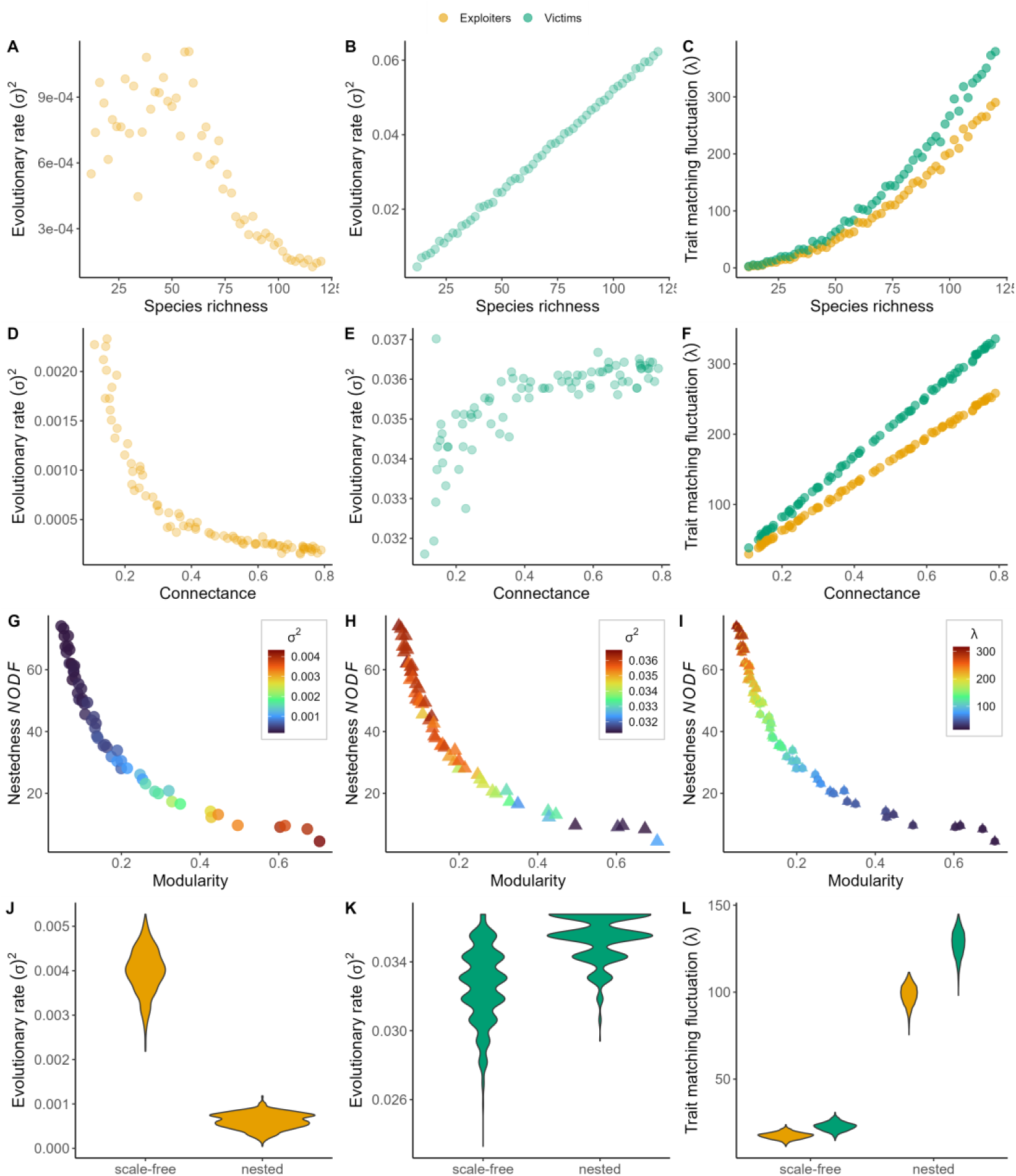
Effect of network structure on species’ evolutionary dynamics in simulated networks. This figure presents results from simulations conducted on artificial networks, where specific network metrics were systematically varied while keeping others constant. Panels A-C show simulations conducted on 86 networks with connectance fixed to the empirical value (C = 0.3). Panels D-F present results from 86 networks with a fixed number of victims and exploiters (30 and 38, respectively, reflecting empirical averages) and varying connectance. Panels G-I display results from 57 networks with empirical richness and connectance but varying modularity. Panels J-L illustrate outcomes from simulations of 100 networks with empirical richness and connectance, grouped by different network distribution types. Each dot represents the mean across 15 replicates per network and across species.

**Figure 2.**
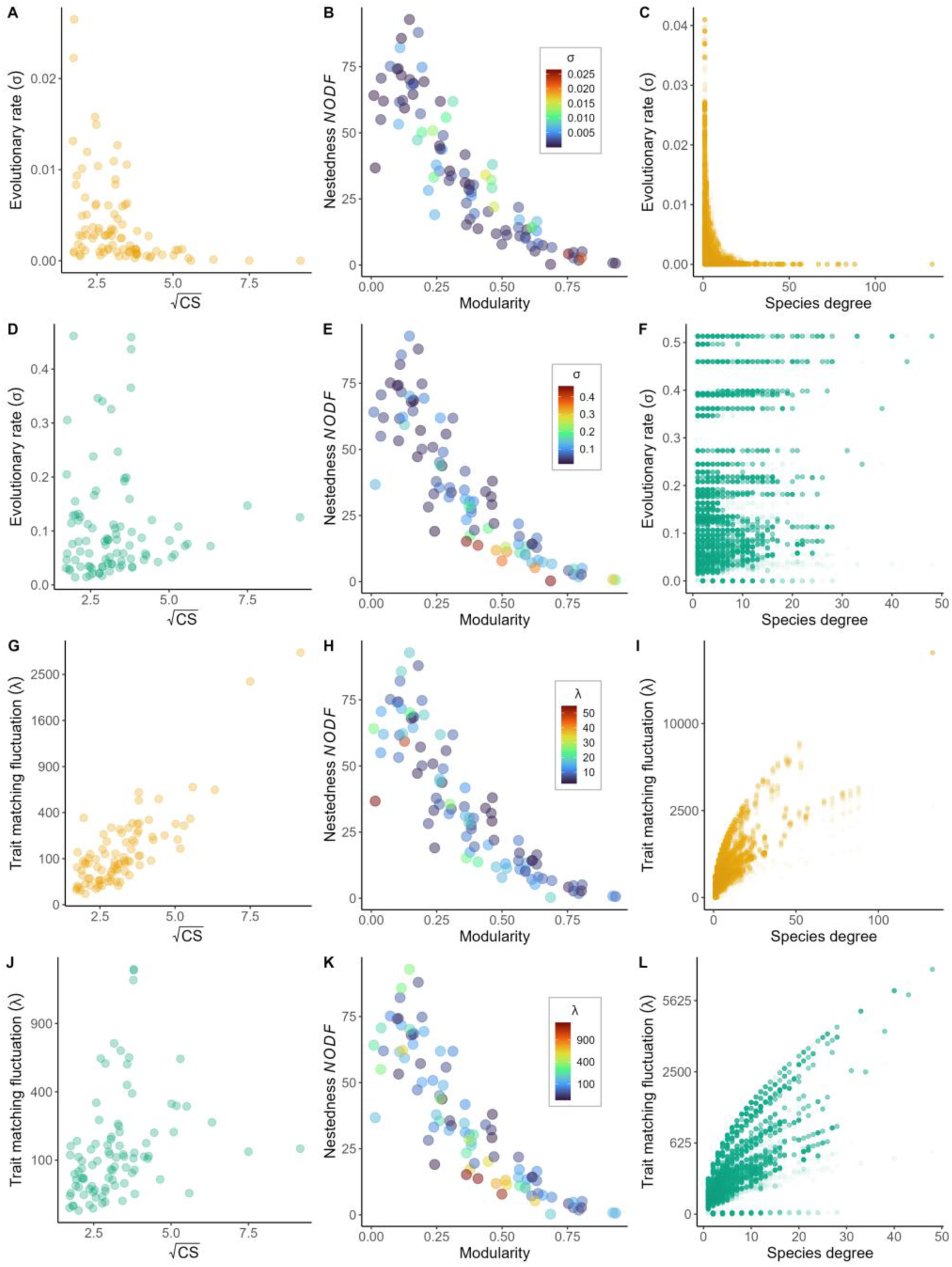
Effect of network structure on evolutionary dynamics in empirical networks. Results from simulations based on 92 empirical network structures. Panels A-C show the evolutionary rate for exploiters, while Panels D-F for victims. Panels G-I illustrate trait matching fluctuations for exploiters, and Panels J-L for victims. Each dot represents the mean across 20 replicates per network and across species.

Trait-matching fluctuations increase with species richness, connectance, and nestedness, revealing a progressive breakdown of coevolution. Simulations with synthetic networks reveal higher trait-matching fluctuations with increasing species richness, connectance, and nestedness, but reduced fluctuations with modularity (Figure 1C,F,I,L). Empirical networks show a similar trend for exploiters, with fluctuations increasing linearly with complexity (Figure 2G). Victims, however, show a hump-shaped relationship, with trait-matching fluctuations peaking at intermediate levels of network complexity (Figure 2J). Fluctuations also increase asymptotically with centrality for both exploiters (Figure 2I). For victims, fluctuations increase gradually at low to moderate centrality, then rise steeply at high centrality, though some low-fluctuation outliers remain (Figure 2L). The relationships with nestedness and modularity are more variable; however, for exploiters, the highest fluctuations in trait matching emerge in less modular structures, whereas for victims, they occur in less nested structures (Figure 2H,K).

Exploiters become more maladapted in complex networks, whereas victims exhibit more stable levels of biotic maladaptation. Exploiters with low values of the complexity index are generally well-adapted to their biotic environment. As network size or species’ degree increases, exploiters become progressively maladapted (Figure 3A,D). Similarly, network load for exploiters rises with both network complexity and species’ degree (Figure 3B,E), indicating greater maladaptation associated with the variance in traits across the network.In contrast, victims show a markedly different relationship. Biotic load is highly variable at low network complexity and for victims with low degrees, spanning a broad range of values. However, as complexity or victim degree increases, this variability diminishes and stabilizes at moderate levels (Figure 3C,F).

**Figure 3.**
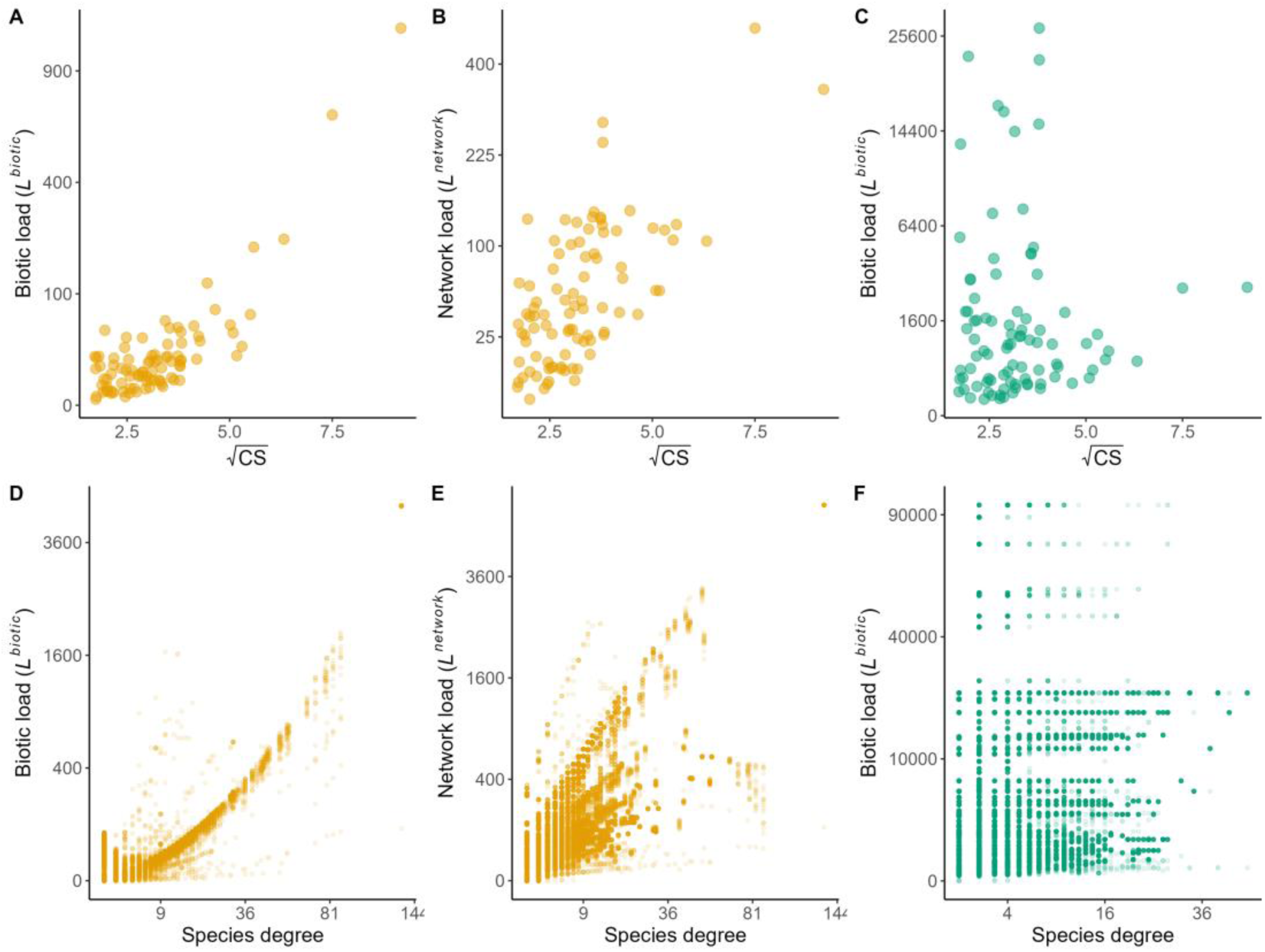
Effect of network structure on biotic adaptation in empirical networks. In these simulations, species began with randomly assigned trait values and evolved in a constant environment until reaching equilibrium. The figure illustrates the relationship between network complexity and various forms of maladaptation: biotic and abiotic loads (Panels A and C) and network load (Panels B and D). Network load represents the incompressible component of maladaptation—independent of the focal species’ trait value—resulting from the variance in trait values among species. The top row displays results for exploiters (A, B), while the bottom row focuses on victims (C, D). Each dot represents the mean across 20 replicates per network and across species. The color legend applies to Panels A and C only.

Biotic evenness (H) is strongly shaped by network structure, with higher evenness emerging in densely connected and nested networks, and lower evenness in modular or scale-free configurations. Biotic evenness increases asymptotically with connectance and decreases almost linearly with modularity (Figure 4A,B). Structures with nested degree distributions exhibit high evenness, whereas scale-free distributions result in low evenness (Figure 4C). Simulations with empirical networks showed similar trends, with evenness being highly variable at low values of the complexity index and species centrality but converging to high values as these increase (Figure 4D,F; see Figure S2 for simulated networks). Evenness also increases with nestedness and decreases with modularity (Figure 4E).

**Figure 4.**
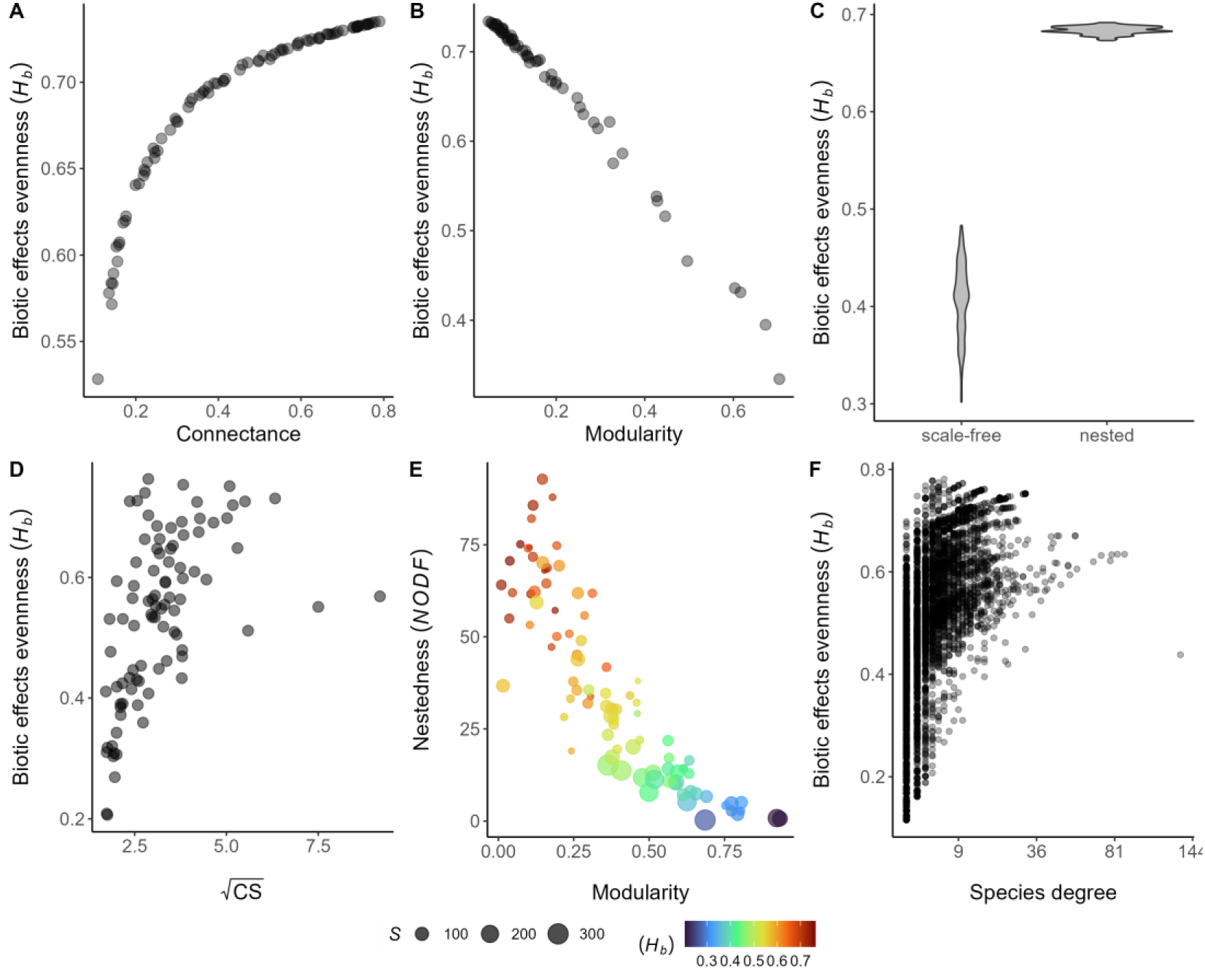
Effect of network structure on species’ perceived biotic effects—simulated and empirical networks. Network structure influences the direct and indirect pathways through which biotic effects affect each species. We measured the evenness of biotic effects (H) perceived by species, where a value of 0 indicates that all effects are concentrated in a single species, while a value of 1 signifies that effects are equally distributed among all species. Panels A-C present the results on simulated networks, where network metrics were systematically varied while controlling for others. Panel A includes 86 networks with a fixed number of victims and exploiters (30 and 38, respectively, reflecting empirical averages) and varying connectance. Panel B shows results from 57 networks with empirical richness and connectance (C = 0.3), but varying modularity. Panel C shows results from 100 networks with empirical richness and connectance, categorized by different network distribution types. Panels D-F display results from simulations using 92 empirical network structures. In panels A,B,D,L, each dot represents the mean across species, and in C and D, the species’ values.

## Discussion

Significant progress has been made in understanding how coevolution shapes network structures (Guimarães et al. 2007, Poisot et al. 2012, Nuismer et al. 2013, Fortuna et al. 2019, de Andreazzi et al. 2020), yet much less is known about how network structure, in turn, influences coevolutionary dynamics and adaptation. Our findings reveal that species embedded in networks with greater interaction diversity and partner overlap, or occupying more central positions, experience a more even distribution of biotic effects across their interactions, affecting their evolutionary dynamics. In our simulations, this structural spread translates into contrasting outcomes between exploiters and victims because we assume different selection rules for each guild: exploiters sum selection across all potential victims, whereas victims escape selection with a trait-distance threshold (Andreazzi et al. 2017, 2020, Pedraza et al. 2025). Biologically, this reflects that consumers can, in principle, adapt to any prey they can attack, while victims are under strong pressure to escape only those exploiters that currently threaten them. As a consequence, structure impacts the guilds differently. Other ways to model selection could change the magnitude of the guild contrast (e.g., applying the threshold to exploiters or using alternative weighting kernels), but our central result remains: network topology predictably affects whether selection is spread or concentrated, and this structural effect shapes evolutionary rates, coevolutionary dynamics, and adaptation.

Exploiters embedded in networks with greater interaction diversity and partner overlap, or those more generalists, experience selection shaped by diverse and often conflicting pressures from multiple victims. These conditions weaken the directional selection required to drive significant trait changes, slowing the directionality of the evolutionary rates and reducing the likelihood of strong coevolutionary feedbacks. These results align with those of de Andreazzi et al. 2017, who found that nested networks sustain fluctuating trait evolution. Conversely, in less connected or more modular networks, or for exploiters specializing in fewer victims, our results show that selection pressures become more concentrated and directional. These focused interactions foster tighter coevolutionary dynamics, resulting in arms race dynamics that drive the rapid spread of advantageous traits, accelerating evolutionary rates (Guimarães et al. 2011). However, our model omits density dependence and trophic regulation, such as exploitative competition, which can reshape fitness outcomes (Holdridge et al. 2016) and potentially counteract spread selection by favoring specialization and strengthening directional selection. Future work should examine how these ecological feedbacks interact with network structure to shape coevolution.

Victims, however, can reduce the number of effective exploiters by evolving their defenses (i.e., differentiating traits enough to escape). This escape mechanism concentrates selection, which can drive rapid trait evolution by allowing victims to optimize their defenses against a smaller subset of closely matched exploiters. However, the variability in victims’ responses in network structures with high interaction diversity and partner overlap suggests that while escape can drive rapid adaptation through aligned selection pressures from different exploiters, conflicting selection pressures from different exploiters may also persist, producing stasis or fluctuating dynamics. In modular or less connected networks, victims interact with fewer exploiters, each exerting distinct and potentially stronger selective pressures, resulting in more variable and localized adaptive responses in the network..

These features of network structure exacerbate exploiters’ difficulty in tracking victims’ evolutionary changes, increasing their maladaptation to the biotic environment. In contrast, victims maintain or even improve their adaptability under high interaction diversity and overlap, benefitting from the more limited responses of exploiters and aligned selective pressures. These findings underscore how more diverse and interlinked networks can disrupt adaptive coevolutionary dynamics, raising questions about its broader implications for community resilience. If these topologies hinder exploiters’ adaptive responses, environmental perturbations may lead to the loss or reconfiguration of interactions, potentially reducing network connectivity and altering its structure. Guimarães et al. (2017) demonstrated that network structures of multiple-partner mutualisms may slow species’ response to environmental changes, and conflicting indirect effects may reduce the magnitude and slow the rate at which species respond to the rapid environmental change in ways that increase their vulnerability to extinction

Furthermore, we found that exploiter’s degree decreases evolutionary rates and increases fluctuating coevolution, making them more maladapted to their victims. This effect also opens new insights on the connection between species’ roles in the network, which is a strong focus for determining keystone species important for ecosystem functioning and guiding conservation strategies (González et al. 2010, Gouveia et al. 2021), and their capacity to evolve and adapt to environmental changes. The evolutionary constraints on generalist exploiters extend beyond their own adaptation, as they are strongly influential in propagating coevolutionary effects through the network influencing the adaptive coevolution of the entire community (Guimarães et al. 2011). Investigating how structural stability and coevolutionary dynamics are balanced in antagonistic networks through central species is thus an additional important future research avenue.

These dynamics may be particularly relevant under scenarios of rapid environmental change, where species within networks can exhibit divergent responses, such as asymmetric range shifts (Carroll et al. 2024). For generalist predators, or species embedded in networks with high interaction diversity and partner overlap, this may reduce their ability to track victims’ responses, potentially disrupting network structure and function. However, our model does not allow for interaction rewiring between species, which is a crucial process in the adaptive dynamics of ecological networks (Raimundo et al. 2018). The structural effects we describe could combine with rewiring and introduce new mechanisms of community-level adaptation. Specifically, spread biotic selection could increase the ability of central species, such as super-generalists, to alternate interacting partners according to the relative defenses of available victims (Thompson 2005, Nuismer & Thomson 2006). This flexibility, together with the enhanced evolutionary stasis observed in central species, could help buffer the network against large structural changes.

We do not know of studies testing this hypothesis, but it may be a common property that confers stability to other biological complex networks, such as in protein regulatory interaction and metabolic networks of yeasts, *Drosophila*, and humans, where highly connected nodes (genes) tend to evolve more slowly than less connected ones (Fraser et al. 2002, Hahn & Kern 2005, Vitkup et al. 2006, Alvarez-Ponce et al. 2017, but see Jovelin & Phillips 2009). Understanding how constraints on central species shape coevolutionary dynamics and network structure over time remains a key but underexplored challenge, which we could start addressing with experiments in microbial communities (Hall et al. 2020).

In natural communities, multiple interaction types coexist. Determining whether our results hold for a broader range of interactions and, more importantly, whether they persist in meta-network communities would be a crucial direction for future research. Pedraza et al. (2024) merged mutualistic and antagonistic coevolving communities and found that the contribution of indirect effects to trait evolution diminishes when increasing the proportion of antagonistic interactions, especially if specialist consumers are antagonists. Moreover, multiple networks can be connected through space, forming spatial meta-networks, which may alter how species are influenced by local structures through the creation of connecting fluxes of biotic effects between networks and species connecting them. Medeiros et al. (2018) found similar results to those we found when analyzing isolated networks, with fewer species and more modular structures favoring higher levels of trait matching than species-rich, nested networks. However, for those connected by gene flow, the effects of environmental selection are canceled out, erasing the conflicting selective pressures on highly interacting species and allowing trait matching and convergence to emerge. These and similar frameworks can explore whether our results translate into a meta-community context.

In addition to not accounting for interaction rewiring, our model makes several simplifying assumptions that, while useful for exploring general relationships, could be relaxed in future work to better reflect biological realism. An important assumption in our model is that each interacting partner contributes equally to the selection gradient acting on a species, reflecting an even distribution of selective pressures. However, if selection on consumer traits instead depends on the relative profit of different prey species, as predicted by optimal foraging theory (Charnov 1976), then consumers would evolve primarily in response to their preferred or most profitable prey, leading to a concentration of selection gradients rather than spread selection. Recent individual-based models show that active consumer choice can strongly shape coevolutionary dynamics, leading to the emergence of trait-convergent modules and stable interaction subgroups over time (de Alvarenga et al. 2022). One way forward to relax the equal-weight assumption is by allowing intuitive within-guild heterogeneity in selection strength—e.g., stronger pressures from obligate than facultative parasites (de Andreazzi et al. 2017).

Another important simplification is our focus on a single trait mediating species interactions. In reality, interactions are often governed by multiple traits, and the direction of trait evolution can shift depending on how these traits interact and co-vary (Nuismer & Doebeli 2004, Assis et al. 2020, Débarre et al. 2014). Using victim-exploiter simulations, Gilman et al. (2012) demonstrated that coevolution in highly dimensional trait spaces, especially under strong trait correlations, can facilitate escape mechanisms for victims. Each additional trait influencing interaction probability provides victims with further opportunities to evolve effective defenses. When traits are highly correlated, victims are more likely to experience facilitated evolution or exploiters constrained adaptation in at least one trait, enabling escape. It is possible that increasing trait dimensionality could produce effects similar to those of network topology, compounding the spread of selection on exploiters and further hampering their adaptation to victims.

Finally, we assume a so-called “magic trait” that simultaneously mediates both biotic and abiotic selection. Although different traits often control these components in realiy, network structures often rely on relatively few traits (Eklöf et al. 2013), and these traits may serve dual roles, influencing both interactions and environmental adaptations (e.g., body size; Martinez 2020) or being genetically correlated with other traits. For example, negative correlations exist between growth and immune response in many organisms (van der Most et al. 2011) or between structural genes in plants and those responsible for mutualistic relationships, such as flower color (Anderson et al. 2014). We expect that the effects described here could be more manifest and relevant in these circumstances.

## Acknowledgements

We thank the Quebec Centre for Biodiversity Science (QCBS) and the Computational Biodiversity Science and Services (BIOS²) training program for supporting AFC’s research stay at the University of São Paulo. Financial support was provided by the NSERC - CREATE Training program in computational biodiversity science and the NSERC Discovery Grant to DG.

