## Supplement 1 for "Trapped in the web: network architectures spread coevolution and shape adaptation"

**Table S1. Parameters and variables used in the coevolutionary model.**

| Notation | Definition | Value |
|---|---|---|
| $z_i(0)$ | Initial quantitative trait value of species *i*. | Sampled from a normal distribution, mean = 0, SD = 0.1 |
| $R$ | Total number of species in the network. | Parametrized from data |
| F | Empirical matrix of species interactions in which $f_{ij}$= 1, if species *i* interacts with *j*, and $f_{ij}$= 0, if *i* does not interact with *j*. | Parametrized from the data |
| $\varphi$ | Evolutionary responsiveness parameter. | 0.25 |
| $S_i$ | Partial selection differential caused by environmental selection. | Variable |
| $M_{ij}$ | Partial selection differential caused by selection imposed by interaction partners. | Variable |
| $\theta_i$ | Trait value of species *i* ($z_i$) favored by stabilizing selection. | $\theta_i = z_i(0)$ |
| $\xi_s$ | Intensity of environmental selection. | 0.5 |

| | | |
|---|---|---|
| $\xi_d$ | Intensity of selection imposed by interaction partners. | 0.5 |
| $p_{ij}$ | Relative preference of exploiter species *i* on victim species *j*. | Variable |
| $b$ | Trait-based interaction bias for exploiters. | 10 |
| $u_{ji}(t)$ | Indicates whether victim species *j* will respond to selection pressure exerted by exploiter *i* at time *t*. | $u_{ji}(t) = \begin{Bmatrix} 1, if \ \lvert z_i - z_j \rvert \ \leq \ \varepsilon \\ 0, if \ \lvert z_i - z_j \rvert > \varepsilon \end{Bmatrix}$ |
| ε | Maximum $\lvert z_{ij}(t) \rvert$ value that induces evolutionary response on victim *j*. | 0.5 |
| $L_i^{biotic}$ | Biotic load for exploiter *i*. | Variable |
| $L_j^{biotic}$ | Biotic load for victim *j*. | Variable |
| $L_i^{network}$ | Network load for exploiter *i*. | Variable |
| P | Matrix of normalized interactions describing the proportional effect of a species *i* in the column on the species *j* in the row. | Variable |
| Q | Weighted P matrix by the amount of biotic selection imposed by biotic interactions ($\xi_d = 0.5$). | Variable |
| T | Matrix of net or total effects. | Variable |
| *H* | Biotic effect's evenness for species *i*, which corresponds to the Shannon index for each row of the *T* matrix. | Variable |
| $\sigma_i$ | Evolutionary rate for species *i*. | Variable |
| $\lambda_{ij}$ | Trait matching fluctuation between species *i* and *j* | Variable |

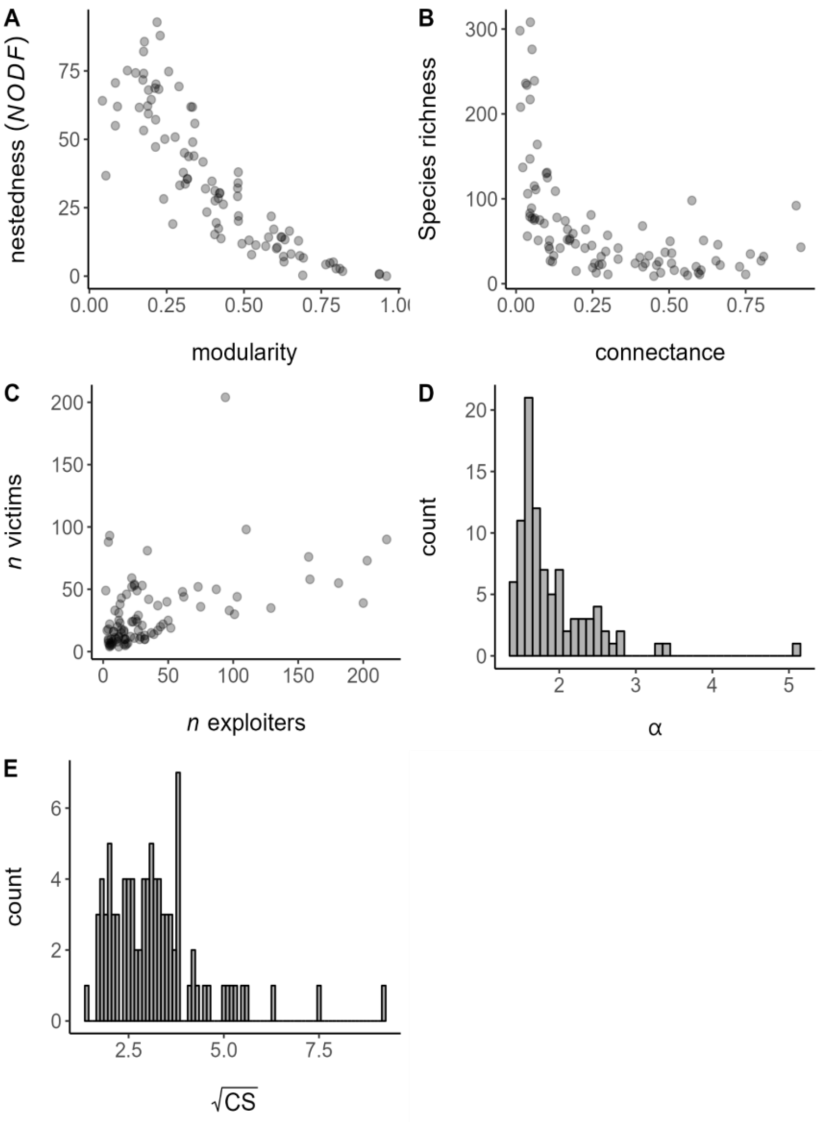


**Figure S1. Structural metrics of the 92 empirical networks used to parametrize the simulations.**

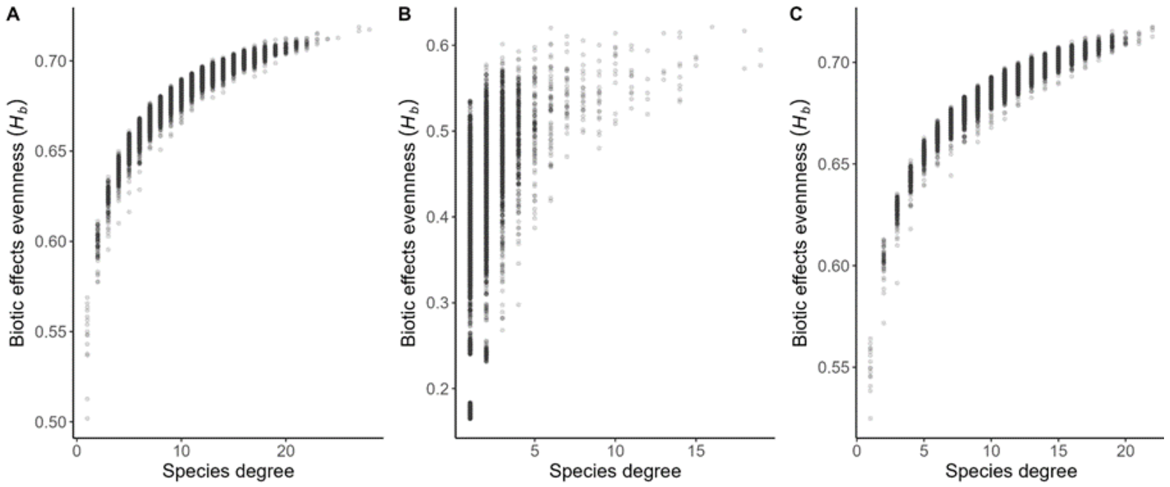


**Figure S2. Evenness of biotic effects and species centrality in simulated networks.**

The plots show the results for sets of 100 networks simulated from random (*A*), scale-free (*B*), and nested (*C*) degree distributions. For all networks, the number of victims and exploiters (30 and 38, respectively) and connectance (0.3) is fixed to the mean of empirical networks. Each dot corresponds to the mean of 15 replicates per network.

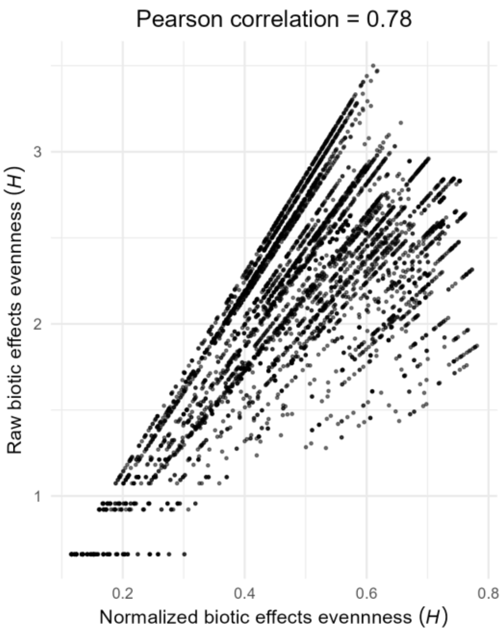

**Figure S3. Comparison of raw and normalized biotic effects evenness (H)—empirical networks.**

The figure depicts the relationship between the raw and normalized biotic effects evenness for all species within 92 empirical network structures, highlighting a strong correlation (Pearson's r = 0.78).

We find a strong correlation (Pearson's r = 0.78) between normalized and raw H (Figure S3.3), with both measures following similar patterns across most network metrics (Figure S3.4). However, some differences emerge in the relation of $H$ with nestedness and modularity, particularly in networks with substantial variation in size. Despite these discrepancies, we used the normalized measure of $H$ in our analyses, as it provides a more consistent basis for comparison across network structures, while offering a clearer interpretation.
